# Comment on “Variation in cancer risk among tissues can be explained by the number of stem cell divisions”

**DOI:** 10.1101/024497

**Authors:** Maxime Tarabichi, Vincent Detours

## Abstract

Tomasetti and Vogelstein (Science 347, 78–81, 2015) claimed that “primary prevention measures are not likely to be very effective” for many cancers because they arise mostly from random mutations fixed during stem cell division, independently of specific genetic or environmental factors. We demonstrate that their calculation for hepatocellular carcinomas overlooked a major subset of tumors proven to be preventable through vaccination. The problem, which is not limited to hepatocellular carcinoma, arises from the general reliance of their analysis on average USA incidences and the omission of incidences in specific risk groups.

---

Tomasetti and Vogelstein (*1*) claimed that for tumors of relatively low incidence arising in organs undergoing many stem cell divisions “primary prevention measures are not likely to be very effective” because they arise mostly from random mutations fixed during stem cell division, independently of specific genetic or environmental factors. This conclusion—which as received much press coverage—has important implications for public health and environmental research and policies.

The authors argued that 2/3 of the variation of cancer incidence among human organs could be explained by the total number of lifetime stem cell divisions (lscd), which, according to them, drives the stochastic accumulation of random mutations. Yet, incidence variation among organs is not informative about incidence variation across different risk groups. For example, worldwide cancer incidence variations and their association with regional risk factors are well documented (*2*). But, the study rests mostly on current average USA incidence statistics and is therefore blind to population-specific risk factors.

Tomasetti and Vogelstein did, however, consider risk-group specific incidences for a few cancers. For example, they calculated the excess risk score (ERS) for hepatocellular cancer (HCC) for the USA subpopulation infected by the hepatitis C virus (HCV) and the non HCV-infected subpopulation. It was 5.36 for HCV and -6.08 in non HCV cancers, which correspond to the D-tumor (deterministic) and R-tumors (replicative) classes, respectively. This seems to support the validity of the ERS. But what would have been the classification of hepatocellular cancer if, as for most other cancers in the study, only the USA average incidence would have been taken into account? The ERS would be -5.65, well within the range of R-tumors, leading to the conclusion that HCC is a less preventable cancer (Fig. 1). This would be a dangerous distraction from the fact that 10 to 33% of them, depending on world regions, are caused by HCV infections that are both preventable and treatable when responsible health policies are implemented. Furthermore, is the non HCV HCC not preventable, as its classification suggests? Fifty nine percents of HCC cases in the developing world are associated with hepatitis virus B infection (*2*), which greatly increases the probability of developing the disease (Fig. 1). Universal HBV vaccination has resulted in a 65-75% reduction of HCC incidence in 6-14 years children from Taiwan (*3*). Other overlooked preventable risk factors for HCC include obesity, alcoholic cirrhosis, exposure to aflatoxin B and schistosomiasis. We focused on HCC due to space constrains, but similar arguments could be made for most cancers analyzed in ref. (*1*).

**Fig. 1.**
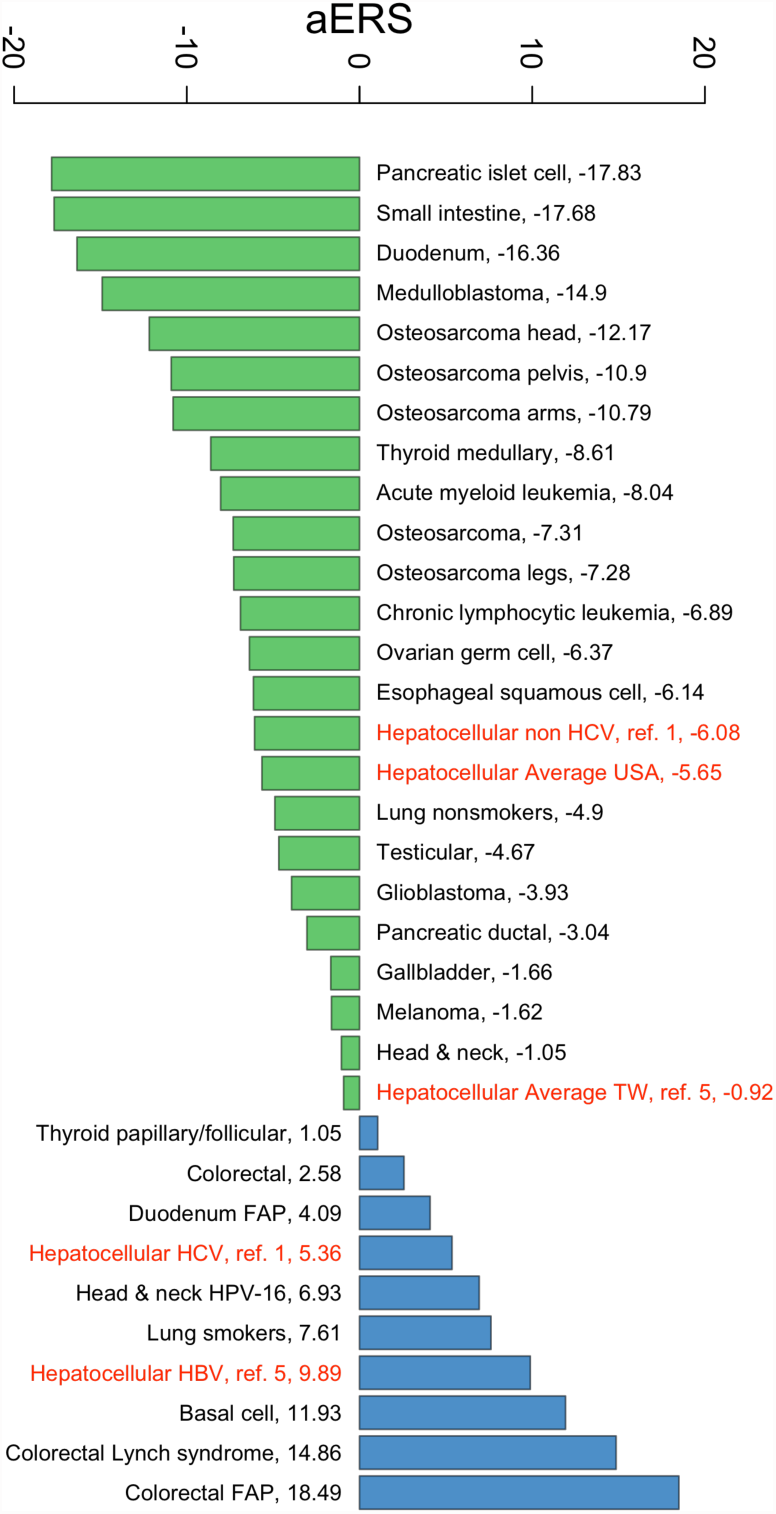
Effect of risk stratification of hepatocellular carcinoma incidence. This figure is an enhanced version of Fig. 2 in ref. (*1*) showing the adjusted ERS (aERS=ERS + 18.49) for human cancers. ‘Hepatocellular Average USA’ and ‘Hepatocellular Average TW’ denotes the entire population of HCC patients, including both HCV and non HCV cases, in the USA and in Taiwan, respectively. Incidence was taken from the SEER database. ‘Hepatocellular HBV’ denotes HCC patients who are also HBV carriers. The HCC lifetime risk for HBsAg-positive patients was taken from ref. (*5*).

We also included in Fig. 1 the ERS for the overall Taiwanese population. It is between the D- and R-tumors and higher than for the USA population. This is consistent with the fact that HCC is more preventable in Taiwan where HCV and HBV are more prevalent and supports, it seems, the potential usefulness of the ERS. Importantly, however, the ERS for all HCC rest on the same lscd estimate, thus incidence data alone would produce the same ranking of the HCC groups [ERS=log_10_(lscd)×log_10_(incidence)]. On a more fundamental level, the ERS does not provide an absolute quantification of determinism because we do not know the baseline ERS for cancers occurring in the *proven* absence of any risk factor. Is this baseline universal or is it organ-specific? If the latter is correct, then the ERS will not be comparable among organs and will not be more informative than incidence data alone, as we have noted for HCC. If not, the ERS scale will be universal and the lscd will add information useful for the comparison of cancer determinism between organs. The modalities of DNA repair varies across the stem cell compartments of different organs (*4*), suggesting an organ-specific baseline.

To our knowledge a substantial variation of the lscd in the general population cannot be excluded. Hence the stratification problem encountered for incidence data may also arise because of lscd variation. The authors wrote that factors “such as those that affect height and weight” could play a role. We are not aware of any relation between cancer and height or weight (disregarding obesity), but we consider highly plausible that tissue repair following chronic and possibly preventable damage may also significantly affect the lscd. Similarly, the relation between mutation rates and lscd is modulated by a range of factors, including DNA repair efficiency and activation of APOBEC DNA mutators (*6*). All of these were averaged out as were most known cancer risk factors.

In order to demonstrate the robustness of the correlation between the lscd and cancer lifetime risk, Tomasetti and Vogelstein varied randomly their lscd estimates over four orders of magnitude. We reproduced this calculation except that incidences were also varied by two orders of magnitude. This calculation confirmed the robustness of the correlation (median ρ=0.5, 95% CI: 0.27-0.66; median p=0.005, 95% CI: 0.00006-0.14). We also collapsed to a single data point cancers sharing the same lscd estimate to address statistical independence concerns. Again, the correlation remained strong (ρ=0.7, p=0.0004).

The remarkable relation between cancer incidence and lscd uncovered by Tomasetti and Vogelstein is statistically robust. The ERS is typically high for known deterministic cancers. But we demonstrated that a cancer with a low ERS may include a sizable fraction of preventable diseases. This proves that their classification scheme, in its current form, is not suitable to gauge the likely effectiveness of prevention measures and to direct funding for research on cancer etiology. Many more risk factors for cancers will likely be discovered in the future. Hence, cancers ascribed to ‘bad luck’ today due to lack of proper risk stratification may someday become explainable and, hopefully, preventable.

